# A new ferrocene derivative blocks KRAS localization and function by oxidative modification at His95

**DOI:** 10.1101/2023.03.28.534499

**Authors:** Kristen M. Rehl, Jayaraman Selvakumar, Don Hoang, Kuppuswamy Arumugam, Alemayehu A. Gorfe, Kwang-jin Cho

**Affiliations:** Department of Biochemistry and Molecular Biology, Boonshoft School of Medicine, Wright State University, Dayton, Ohio 45435, USA; Department of Chemistry, College of Science and Mathematics, Wright State University, Dayton, Ohio 45435, USA; Department of Integrative Biology and Pharmacology, McGovern Medical School, University of Texas Health Science Center, Houston, Texas 77030, USA

**Keywords:** Ferrocene, KRAS, Oxidation, Plasma membrane, ROS

## Abstract

Ras proteins are membrane-bound GTPases that regulate essential cellular processes at the plasma membrane (PM). Constitutively active mutations of K-Ras, one of the three Ras isoforms in mammalian cells, are frequently found in human cancers. Ferrocene derivatives, which elevate cellular reactive oxygen species (ROS), have shown to block the growth of non-small cell lung cancers (NSCLCs) harboring oncogenic mutant K-Ras. Here, we developed and tested a novel ferrocene derivative on the growth of human pancreatic ductal adenocarcinoma (PDAC) and NSCLC. Our compound inhibited the growth of K-Ras-dependent PDAC and NSCLC and abrogated the PM binding and signaling of K-Ras, but not other Ras isoforms. These effects were reversed upon antioxidant supplementation, suggesting a ROS-mediated mechanism. We further identified K-Ras His95 residue in the G-domain as being involved in the ferrocene-induced K-Ras PM dissociation via oxidative modification. Together, our studies demonstrate that the redox system directly regulates K-Ras PM binding and signaling via oxidative modification at the His95, and proposes a role of oncogenic mutant K-Ras in the recently described antioxidant-induced metastasis in K-Ras-driven lung cancers.

## Introduction

Ras proteins are a family of small GTPases primarily localized to the plasma membrane (PM) and function as molecular switches, cycling between a GDP-bound inactive state and a GTP-bound active state (Chen *et al*, 2021; McCormick, 2016). In response to activation by upstream receptor kinases, guanine exchange factors (GEFs) bind to Ras to facilitate the release of GDP and binding of GTP, which confers Ras the active conformation for binding its downstream effectors (Chen *et al*., 2021; McCormick, 2016). Ras proteins are involved in a variety of cellular pathways that regulate cell differentiation, proliferation and survival (Chen *et al*., 2021; Jancik *et al*, 2010). GTPase-activating proteins (GAPs) promote Ras GTPase activity, enhancing Ras-induced GTP hydrolysis to GDP (Chen *et al*., 2021; McCormick, 2016). There are three Ras isoforms that are ubiquitously expressed in mammalian cells: K-, H- and N-Ras. K- Ras undergoes alternative splicing on the fourth exon, yielding K-Ras4A and K-Ras4B, where K-Ras4B, hereafter K-Ras, is the predominant K-Ras protein expressed in mammalian cells (Cox *et al*, 2014). Constitutively active mutations of Ras are found in ∼19% of all human cancers, with ∼75% of those being K-Ras (Prior *et al*, 2020). Oncogenic mutant K-Ras are found in ∼88% pancreatic, ∼50% colorectal and ∼30% of lung cancer (Prior *et al*., 2020), and a group of newly developed K-Ras G12C direct inhibitors have shown promise in clinical trials. Sotorasib and adagrasib are FDA-approved small molecules that directly bind to the GDP-bound K-Ras G12C mutant and form a covalent bond to the mutant Cys, which locks K-Ras in the inactive conformation, thereby blocking its signaling (Canon *et al*, 2019; Hallin *et al*, 2020; Ostrem *et al*, 2013). While these inhibitors exhibit pronounced anti-cancer effects in K-Ras G12C tumor mice models and in clinical trials (Canon *et al*., 2019; Hallin *et al*., 2020), the K-Ras G12C mutation is found only in ∼3% pancreatic, ∼4% colorectal and ∼13% of lung cancers that harbor any oncogenic mutations in K-Ras, making it effective against only a small subset of the K-Ras-driven cancers (Cox *et al*., 2014; Prior *et al*., 2020).

One of the strategies for blocking pan-oncogenic mutant K-Ras activity is to dissociate it from the PM. Experimental data show that K-Ras, regardless of mutation status, activates its downstream effectors primarily at the inner PM leaflet, and that dissociating K-Ras from the PM blocks K-Ras signal output and the growth of K-Ras-driven cancers (Cho *et al*, 2016a; Cho *et al*, 2012b; Cho *et al*, 2016b; Garrido *et al*, 2020; Kattan *et al*, 2019; Kattan *et al*, 2021; Kovar *et al*, 2020; Miller *et al*, 2019; Tan *et al*, 2019; van der Hoeven *et al*, 2013). Ras proteins have two targeting signals for stable interaction with the PM. The first signal is the C-terminal CAAX motif that undergoes a series of post-translational modifications to generate a farnesyl-cysteine-methyl-ester, allowing Ras binding to cellular membranes (Apolloni *et al*, 2000; Prior & Hancock, 2012). For H- and N-Ras, the second targeting signal is palmitoyl moieties adjacent to the farnesylated Cys, conferring their stable PM binding. The second targeting signal of K-Ras is its C-terminal polybasic domain, a stretch of six lysine residues near the farnesyl moiety, which forms favorable electrostatic interactions with phosphatidylserine (PtdSer), an anionic phospholipid enriched in the PM inner leaflet (Hancock *et al*, 1990; Yeung *et al*, 2008; Zhou *et al*, 2017). Recent studies have identified three molecular mechanisms that regulate K-Ras transport to, and interaction with the PM: ***(i)*** disrupting K-Ras binding to its chaperone protein, phosphodiesterase 6 delta (PDE6δ), ***(ii)*** phosphorylating K-Ras at Ser181 and ***(iii)*** reducing PM PtdSer content (Gorfe & Cho, 2019; Henkels *et al*, 2021). Under basal conditions, H-, N- and K- Ras proteins are farnesylated, and PDE6δ binds endocytosed Ras proteins via the farnesyl moiety and returns them to the PM. Disrupting this interaction redistributes all three Ras isoforms to endomembranes (Chandra *et al*, 2012; Schmick *et al*, 2014; Zimmermann *et al*, 2013). Also, protein kinase C and G directly phosphorylate K-Ras, but not other Ras isoforms, at Ser181 via independent mechanisms. This perturbs K-Ras PM binding and mislocalizes K-Ras from the PM to endomembranes (Bivona *et al*, 2006; Cho *et al*., 2016a; Kovar *et al*., 2020). The other regulator for K-Ras PM localization is PtdSer enrichment in the inner PM leaflet. Depletion of PM PtdSer by perturbing mechanisms that maintain PM PtdSer abundance dissociates K-Ras from the PM and blocks K-Ras signal output and the growth of human cancers harboring oncogenic mutant K-Ras *in vitro* and *in vivo* (Cho *et al*., 2012b; Cho *et al*., 2016b; Kattan *et al*., 2021; van der Hoeven *et al*, 2018).

Ferrocene derivatives encompass a diverse range of compounds, but they all contain the ferrocene molecule, an iron atom (Fe) sandwiched between two cyclopentadienyl ligands (Gasser *et al*, 2011; Kealy & Pauson, 1951; Wilkinson *et al*, 1952). The ferrocene molecule is insoluble and inactive, requiring additional functional groups to enter the cell (Fayaz Ali Larik, 2016; Peter & Aderibigbe, 2019). Once inside, it undergoes a one-electron oxidation, Fe^2+^ to Fe^3+^, to form the active ferrocenium cation (Fayaz Ali Larik, 2016; Gasser *et al*., 2011). The general mechanism of ferrocene derivatives involves the elevation of cellular reactive oxygen species (ROS) to induce cellular damage and apoptosis (Melendez, 2012; Mojzisova *et al*, 2014). For example, a ferrocene conjugated to a gold(I)N-heterocyclic carbene complexes elevates cellular ROS by irreversibly binding to and inhibiting thioredoxin reductase, an essential component of the thioredoxin antioxidant system (Arambula *et al*, 2016). Human cancers harboring oncogenic mutant K-Ras are susceptible to changes in cellular ROS due to their metabolic reprogramming, and ROS-elevating agents are effective in inhibiting their growth (Arambula *et al*., 2016; Foo & Pervaiz, 2019; McCall *et al*, 2017). Also, ferrocene derivatives exhibit anti-cancer effects in lung cancer cells harboring oncogenic mutant K-Ras (Arambula *et al*., 2016; Domarle *et al*, 1998; Perez *et al*, 2015; Peter & Aderibigbe, 2019). Thus, ferrocene derivatives may be used for targeting K-Ras-driven cancers.

In this study, we developed a novel ferrocene complex, C_16_H_20_FeClNO, which inhibits the growth of K-Ras-driven human pancreatic ductal adenocarcinoma (PDAC) and non-small cell lung cancer (NSCLC) cells. We also discovered that elevation of cellular ROS disrupts the PM binding and signaling of K-Ras, but not other Ras isoforms, through oxidative modification of K-Ras His95 residue. Moreover, our findings may provide an additional role of oncogenic mutant K-Ras in the recently reported antioxidant-induced metastasis of K-Ras-driven NSCLC (Lignitto *et al*, 2019; Wiel *et al*, 2019).

## Results and Discussion

### Synthesis of C_16_H_20_FeClNO

Compound **1** was prepared by treating acetylferrocene with a premixed mixture of commercially available materials, bis(dimethylamino)methane, phosphoric, and acetic anhydride (Fig. 1A). After workup, compound **1** was obtained as a crude product, which was purified using column chromatography to yield the pure product as orange-yellow oil in 71% yield. The obtained product was characterized using ^1^H and ^13^C NMR spectroscopy to confirm the identity of compound **1** (Fig. EV1A and B). The resulting oil was further treated with HCl dissolved in diethyl ether to yield the pure compound **2** in 95% yield (Fig. 1A). The obtained product was subjected to ^1^H and ^13^C NMR spectroscopy, and the identity was confirmed using high-resolution mass spectroscopy and elemental analysis (Figs. EV1C and D). The obtained compound **2** (C_16_H_20_FeClNO) was further subjected to biological studies.

**Figure 1.**
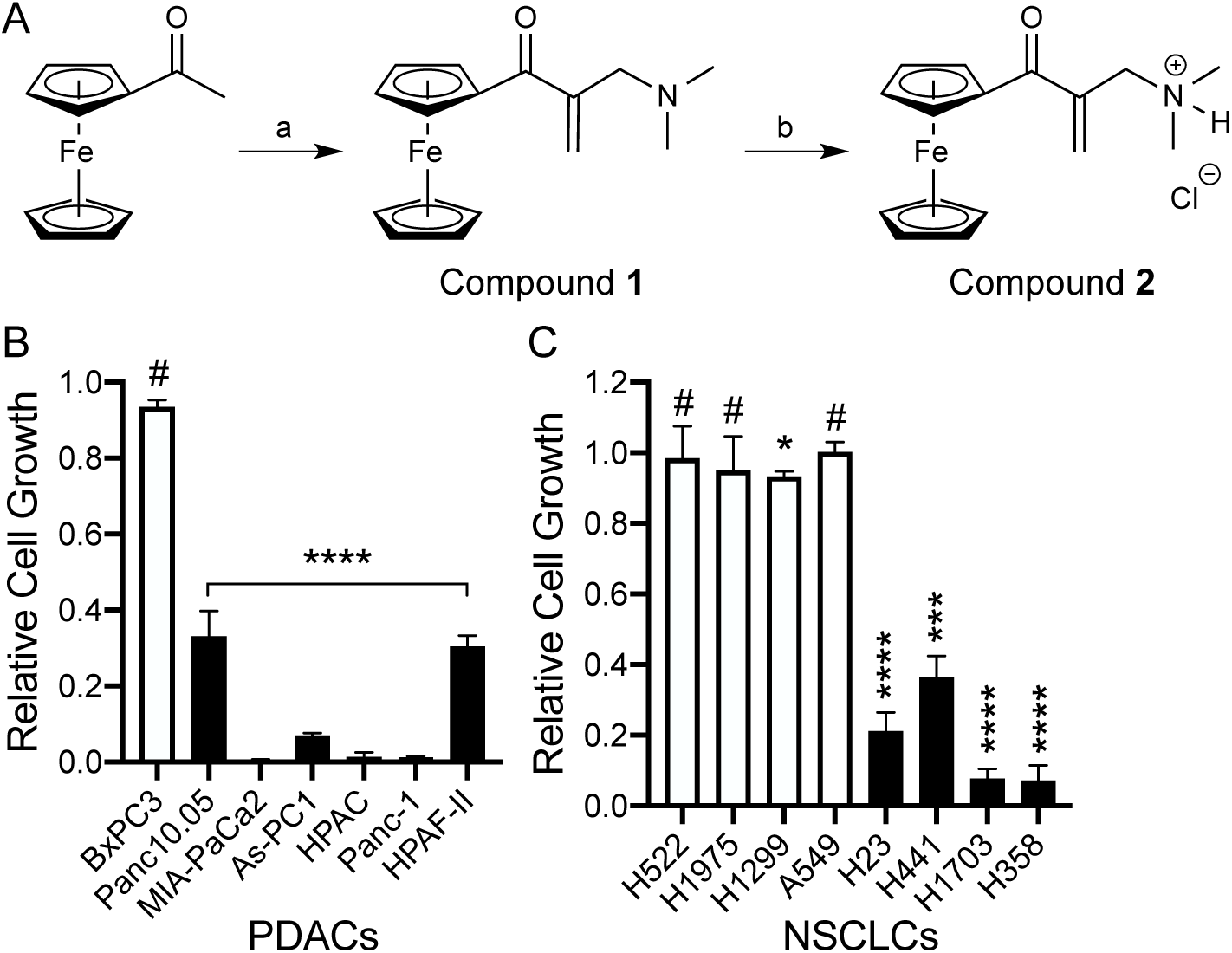
C_16_H_20_FeClNO blocks the growth of K-Ras-dependent human PDAC and NSCLC cells. (**A**) Systematic synthesis of ferrocene derivatives. **a** = CH_2_(N(CH_3_)_2_)_2_, H_3_PO_4_, CH_3_COOH, 110 °C, 4 h; **b** = HCl in O(CH_2_CH_3_)_2_, CH_2_Cl_2_, 0 °C - 23 °C. A panel of pancreatic ductal adenocarcinoma (PDAC) (**B**) and non-small cell lung cancer (NSCLC) cells (**C**) were plated on 96-well plates and treated with 50 μM C_16_H_20_FeClNO for 3 days. Complete growth medium with the compound was replaced every 24h. Cell proliferation was analyzed using CyQuant proliferation assay. The graphs show the mean cell proliferation ± S.E.M. from three independent experiments relative to that for control cells (DMSO-treated). Open and closed bars represent cancer cells, in which their growth is K-Ras-independent and - dependent, respectively. Significant differences between control (DMSO-treated) and C_16_H_20_FeClNO-treated cells were assessed using Student’s *t*-tests (* p<0.05, *** p<0.001, **** p<0.0001, # not significant).

### C_16_H_20_FeClNO inhibits the growth of K-Ras-dependent human cancers

A wide range of human cancers harboring oncogenic mutant K-Ras reprogram their signaling network so that their survival and growth depend on the oncogenic K-Ras signaling, a phenomenon called K-Ras addiction or dependence (Hayes *et al*, 2016; Singh *et al*, 2009; Singh *et al*, 2012; Weinstein & Joe, 2008). Since ferrocene derivatives have shown to inhibit the growth of lung cancer cells that harbor oncogenic mutant K-Ras (Arambula *et al*., 2016; Fayaz Ali Larik, 2016; Wang *et al*, 2020), we examined if our ferrocene compound, C_16_H_20_FeClNO, inhibits the growth of K-Ras-dependent human cancer cell lines. A panel of human PDAC and NSCLC cells that are K-Ras-dependent or -independent were grown in the presence of C_16_H_20_FeClNO, and cell proliferation was measured. Our data show that the compound significantly reduced the proliferation of K-Ras-dependent PDAC cell lines by 70-90%, but not BxPC-3, a K-Ras-independent PDAC cell line harboring WT K-Ras (Fig. 1B). C_16_H_20_FeClNO also blocked the growth of K-Ras-dependent NSCLC cell lines by 60-90%, but had minimal effects on K-Ras-independent cell lines (Fig. 1C). These data suggest that C_16_H_20_FeClNO specifically inhibits the growth of K-Ras-dependent human PDAC and NSCLC cells.

### C_16_H_20_FeClNO translocates K-Ras from the plasma membrane to endomembranes

K-Ras conducts signal transduction by recruiting its downstream effectors primarily to the PM, and blocking the PM localization of K-Ras abrogates K-Ras signaling and the growth of K-Ras-dependent cancers (Cho *et al*., 2016a; Cho *et al*., 2012b; Cho *et al*., 2016b; Garrido *et al*., 2020; Kovar *et al*., 2020; Miller *et al*., 2019; van der Hoeven *et al*., 2018). To examine if C_16_H_20_FeClNO perturbs K-Ras PM binding, we directly measured K-Ras PM binding by quantitative electron microscopy (EM). Intact basal PM sheets from Madin-Darby canine kidney (MDCK) cells stably expressing GFP-tagged oncogenic mutant K-Ras (K-RasG12V) or -H-RasG12V and treated with the compound for 48h were labeled with anti-GFP antibody conjugated to 4.5-nm gold particles and imaged by EM (Prior *et al*, 2001; Prior *et al*, 2003). Our data show that the total number of gold particles for GFP-K-RasG12V, but not -H-RasG12V, was significantly decreased after C_16_H_20_FeClNO treatment (Figs. 2A and EV2). Ras proteins in the PM are spatially organized into nano-scale domains, called nanoclusters, that are essential for high-fidelity Ras signal transduction (Cho *et al*, 2012a; Prior *et al*., 2003; Tian *et al*, 2007). Thus, we examined the spatial organization of Ras proteins remaining at the PM after C_16_H_20_FeClNO treatment. Our data show a significant reduction in the nanoclustering of K-RasG12V, but not H-RasG12V after C_16_H_20_FeClNO treatment (Fig. 2B), suggesting that C_16_H_20_FeClNO disrupts the PM binding and nanoclustering of K-Ras, but not H-Ras.

**Figure 2.**
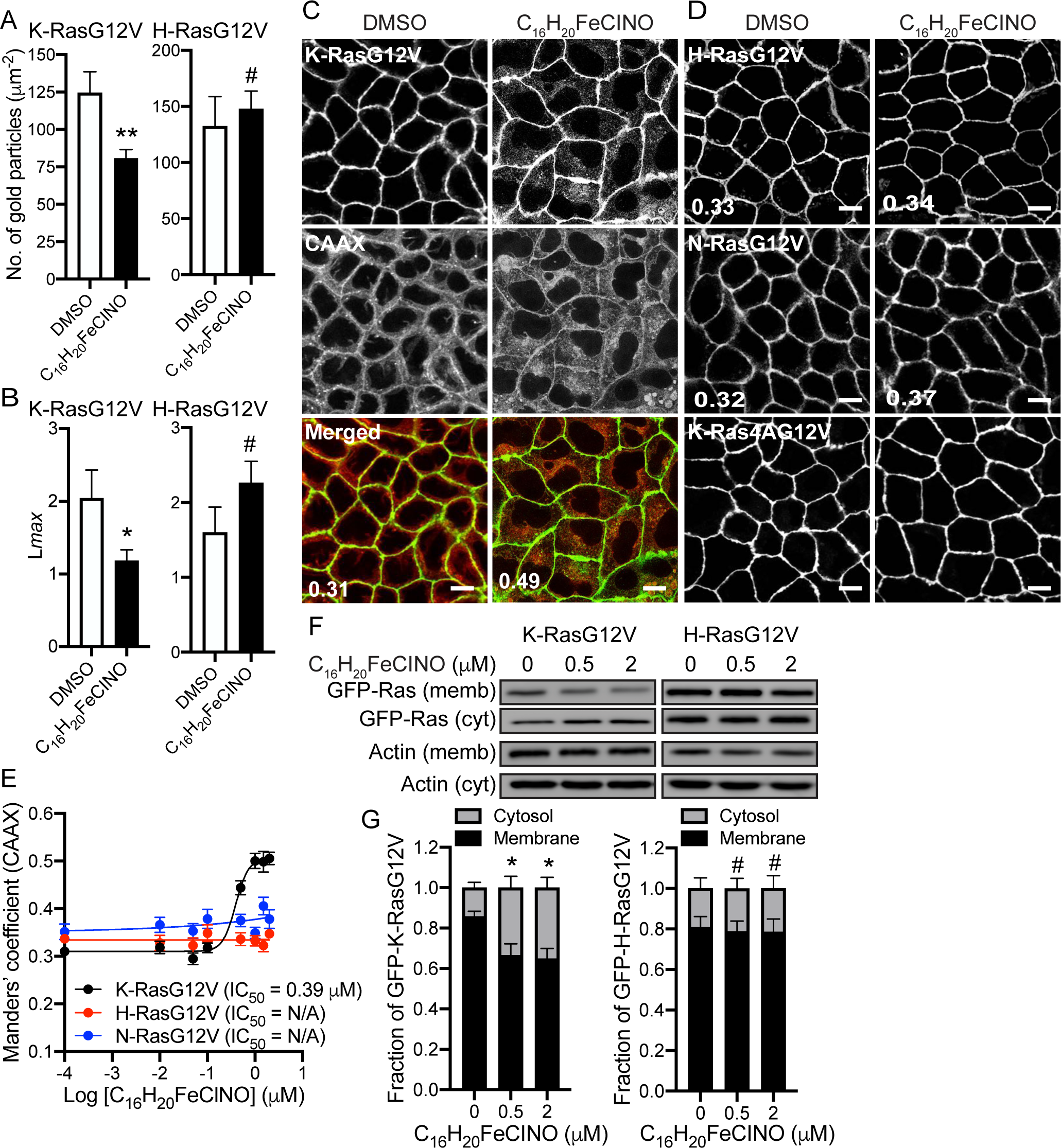
C_16_H_20_FeClNO translocates K-Ras from the PM to the cytosol and endomembranes. (**A**) Basal PM sheets prepared from MDCK cells expressing GFP-K-RasG12V or -H-RasG12V and treated with 2 μM C_16_H_20_FeClNO for 48 h were labeled with anti-GFP-conjugated gold and visualized by EM. Representative EM images are shown in (Fig. **EV2)**. The graphs show a mean number of gold particles ± S.E.M (n ≥ 15). Significant differences between control (DMSO-treated) and C_16_H_20_FeClNO-treated cells were assessed by Student’s *t*-tests. (**B**) Spatial mapping of the same gold-labeled PM sheets was performed. The peak values, *L*_max_, of the respective weighted mean K-function *L(r) - r* curves are shown as bar graphs (*n* ≥ 15). Significant differences between control (DMSO-treated) and C_16_H_20_FeClNO-treated cells were evaluated with bootstrap tests (* p<0.05, ** p<0.01, # - not significant). MDCK cells stably co-expressing mCherry-CAAX and (**C**) GFP-K-RasG12V, (**D**) -H-RasG12V or -N-RasG12V, or expressing GFP-K-Ras4A G12V only were treated with various concentrations of C_16_H_20_FeClNO for 48 h. Cells were fixed with 4% PFA and imaged by confocal microscopy. Representative images of 2 μM C_16_H_20_FeClNO-treated cells are shown. Inserted values represent an estimated mean fraction of mCherry-CAAX co-localizing with GFP-K-RasG12V, -H-RasG12V or -N-RasG12V calculated by Manders’ coefficient from three independent experiments. Scale bar: 10 μm. (**E**) IC_50_s were estimated from the dose-response plots. (**F**) MDCK cells stably expressing GFP-K-RasG12V or -H-RasG12V were treated with C_16_H_20_FeClNO for 48 h. Cell lysates were fractionated into membrane (memb.) and cytosol (cyt.) fractions, and RasG12V levels were measured by immunoblotting using an anti-GFP antibody. Representative blots from three independent experiments are shown. Actin blots were used as loading controls. (**G**) The graphs show the mean membrane-bound fraction ± S.E.M of RasG12V, calculated as membrane/(membrane + cytosol) from three independent experiments. Significant differences of membrane fraction between control (DMSO-treated) and C_16_H_20_FeClNO-treated cells were assessed by one-way ANOVA tests (* p<0.05, # - not significant).

To further investigate the cellular localization of K-RasG12V after C_16_H_20_FeClNO treatment, MDCK cells stably co-expressing mCherry-tagged CAAX, a generic endomembrane marker (Cho *et al*., 2012b; Choy *et al*, 1999), and GFP-K-RasG12V were treated with various concentrations of C_16_H_20_FeClNO for 48h and imaged by confocal microscopy. To quantitate the extent of K-Ras translocation from the PM to endomembranes, an IC_50_ value for Ras PM dissociation was derived from Manders’ coefficient, which calculates the fraction of mCherry-CAAX co-localized with GFP-K-RasG12V (Cho *et al*., 2012b; Manders *et al*, 1993). Our data show that C_16_H_20_FeClNO translocated K-RasG12V to endomembranes with an IC_50_ of 0.39 μM (Figs. 2C and E). Our confocal microscopy further shows the compound did not disrupt the PM localization of GFP-H-RasG12V, -N-RasG12V and -K-Ras4A G12V (Figs. 2D and E). Cellular fractionation assay further reveals that the membrane-associating K-RasG12V is reduced while the cytosolic K-RasG12V is elevated after C_16_H_20_FeClNO treatment. No changes were observed with H-RasG12V (Figs. 2F and G). These data suggest C_16_H_20_FeClNO selectively translocates K-Ras from the PM to the cytosol and endomembranes.

To examine to which cellular organelles the mislocalized K-Ras is redistributed, we co-stained cells expressing GFP-K-RasG12V with organelle markers after C_16_H_20_FeClNO treatment. Our data show that K-RasG12V co-localized with Rab5A-, Rab7A-, LAMP1-, GALNT1-, and ER-tracker-positive structures after the treatment (Figs. EV3A – D and F), suggesting that C_16_H_20_FeClNO translocates K-Ras to the early endosome, late endosome, lysosome, the Golgi complex, and ER. K-RasG12V did not co-localize with a mitochondrial protein, PDHA1, indicating K-RasG12V did not translocate to mitochondria (Fig. EV3E). Together, these data suggest that C_16_H_20_FeClNO disrupts K-Ras/PM binding and nanoclustering of the remaining K-Ras at the PM, but not other Ras isoforms, and that the mislocalized K-Ras translocates to the cytosol and various cellular organelles except mitochondria.

### C_16_H_20_FeClNO blocks the K-Ras/MAPK signaling

To investigate if the disrupted K-Ras PM binding and nanoclustering after C_16_H_20_FeClNO treatment translates to Ras signaling, MDCK cells expressing GFP-tagged different oncogenic mutant Ras isoforms were treated with C_16_H_20_FeClNO and probed for phosphorylated ERK (ppERK) and Akt (S473), the two most well-studied Ras downstream effectors. Our data show that C_16_H_20_FeClNO significantly decreased the level of ppERK, but not pAkt, in a dose-dependent manner only in K-RasG12V-expressing cells (Figs. 3A and B). We also measured total K-Ras protein expression after C_16_H_20_FeClNO treatment since K-Ras/PM dissociation by different mechanisms differentially regulate K-Ras protein expression (Cho *et al*., 2012b; Garrido *et al*., 2020; van der Hoeven *et al*., 2013). Our data show that the compound did not change the protein expression of Ras isoforms (Figs. 3A and B). A previous study reported that K-Ras is a more potent activator of the Raf-1/MAPK signaling than H-Ras, while H-Ras is a more potent activator of the PI3K/Akt signaling (Yan *et al*, 1998). Thus, it is plausible that C_16_H_20_FeClNO blocks the PM binding and nanoclustering of K-Ras, but not other Ras isoforms, resulting in significant inhibition of the Raf-1/MAPK signaling with minimal effects on the PI3K/Akt signaling. Moreover, C_16_H_20_FeClNO did not induce cleavage of caspase-3 and PARP-1 3, suggesting the compound does not induce apoptosis at the concentration that blocks K-Ras/PM binding and K-Ras/MAPK signaling (Fig. EV4). Together with Fig. 2, our data suggest that C_16_H_20_FeClNO inhibits K-Ras PM binding, nanoclustering and K-Ras/MAPK signaling.

**Figure 3.**
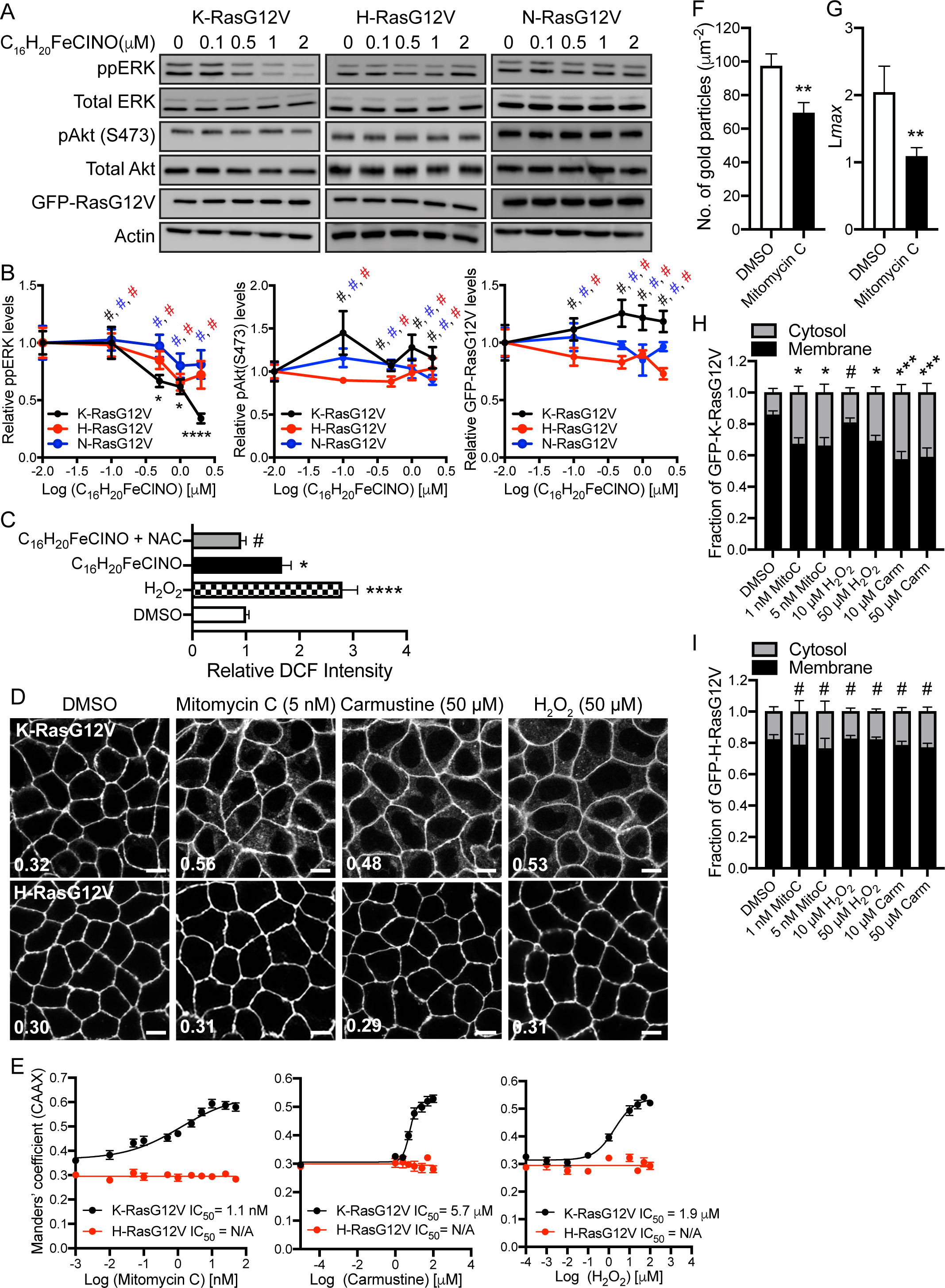
C_16_H_20_FeClNO blocks the K-Ras/MAPK signaling, and ROS-elevating agents dissociates K-Ras from the PM. (**A**) MDCK cells stably expressing GFP-K-RasG12V, -H-RasG12V or -N-RasG12V were treated with C_16_H_20_FeClNO for 48 h. Cell lysates were immunoblotted for phosphorylated ERK (ppERK), Akt (S473) and GFP-RasG12V. Actin, total ERK and Akt blots are shown as loading controls. Representative blots from three independent experiments are shown. (**B**) The graphs show the relative mean of ppERK, pAkt (S473) and GFP-RasG12V ± S.E.M. from three independent experiments. (**C**) WT MDCK cells were seeded onto 96-well plates and grown in the presence 25 μM DCFH_2_-DA with H_2_O_2_ (100 μM), C_16_H_20_FeClNO (2 μM) and/or NAC (500 μM) for 48h. DCF, a readout for cellular ROS was measured by a fluorescent plate reader using emission = 495 nm and excitation = 527 nm. (**D**) MDCK cells co-expressing mCherry-CAAX and GFP-K-RasG12V or -H-RasG12V were treated with various concentrations of drugs that elevate cellular ROS for 48 h, and imaged by confocal microscopy. Representative images of GFP-RasG12V are shown. Inserted values represent an estimated mean fraction of mCherry-CAAX co-localized with GFP-RasG12V calculated by Manders’ coefficient from three independent experiments. Scale bar: 10 μm. (**E**) IC_50_s was estimated from the dose-response plots. (**F**) Basal PM sheets prepared from MDCK cells expressing GFP-K-RasG12V and treated with 5 nM mitomycin C for 48h were labeled with anti-GFP-conjugated gold and visualized by EM. The graph show a mean number of gold particles ± S.E.M (n ≥ 15). (**G**) Spatial mapping of the same gold-labeled PM sheets was performed. The peak values, *L*_max_, of the respective weighted mean K-function *L(r) - r* curves are shown as bar graphs (*n* ≥ 15). (**H**) MDCK cells stably expressing GFP-K-RasG12V or (**I**) -H-RasG12V were treated with indicated drugs for 48 h. Cell lysates were fractionated into membrane and cytosol fractions, and RasG12V levels were measured by immunoblotting using an anti-GFP antibody. The graphs show the mean membrane-bound fraction ± S.E.M of RasG12V, calculated as membrane/(membrane + cytosol) from three independent experiments. Significant differences between control (DMSO-treated) and drug-treated cells were evaluated by one-way ANOVA tests for (**B, C, H** and **I**), Student’s *t*-tests for (**F**), and bootstrap tests for (**G**) (* p<0.05, ** p<0.01, *** p<0.001, **** p<0.0001, # - not significant).

### C_16_H_20_FeClNO disrupts K-Ras PM binding and signal output via elevating cellular ROS

Since ferrocene derivatives elevate cellular ROS levels (Acevedo-Morantes *et al*, 2012; Arambula *et al*., 2016), we examined if our compound elevates cellular ROS. Briefly, WT MDCK cells were treated with C_16_H_20_FeClNO in the presence of DCFH_2_-DA (2′,7′- dichlorodihydrofluorescein diacetate), a chemically reduced form of fluorescein. Once it enters a cell, esterases convert it to DCFH, which is further metabolized to highly fluorescent DCF upon exposure to cellular ROS, a readout for cellular ROS levels (Acevedo-Morantes *et al*., 2012). Our data show that C_16_H_20_FeClNO increased DCF production while co-treatment with N-acetylcysteine (NAC), a general antioxidant (Spagnuolo *et al*, 2006) reversed it (Fig. 3C), confirming that C_16_H_20_FeClNO elevates cellular ROS at the concentration that blocks K-Ras/PM binding and K-Ras signaling. To validate the role of cellular ROS in K-Ras/PM binding, we tested other chemical modulators that elevate cellular ROS by blocking the glutathione or thioredoxin antioxidant systems. MDCK cells co-expressing mCherry-CAAX and GFP-K-RasG12V or -H-RasG12V were treated with mitomycin C, a thioredoxin reductase inhibitor, carmustine, a glutathione reductase inhibitor, or hydrogen peroxide (H_2_O_2_) (An *et al*, 2011; Paz *et al*, 2012; Yokoyama *et al*, 2017), and cellular localization of Ras proteins were imaged. Our data show that these compounds mislocalized K-RasG12V, but not H-RasG12V (Figs. 3D and E). EM analysis further shows that mitomycin C blocks K-Ras/PM binding, nanoclustering and K-Ras/MAPK signaling (Figs. 3F - G and EV5). Moreover, subcellular fraction assay reveals that mitomycin C, carmustine and a high concentration of H_2_O_2_ significantly increased the cytosolic fraction of GFP-K-RasG12V, but not H-RasG12V (Figs. 3H and I). These data suggest that cellular ROS disrupts the PM binding and signaling of K-Ras.

To validate our data, we repeated these experiments after NAC supplementation. MDCK cells expressing GFP-K-RasG12V were co-treated with NAC and C_16_H_20_FeClNO, mitomycin C or H_2_O_2_, and cellular localization of K-RasG12V was imaged. Our confocal and electron microscopy show that NAC supplementation restored K-RasG12V PM binding and nanoclustering (Figs. 4A – D). Moreover, our immunoblot data show that co-treatment of C_16_H_20_FeClNO with NAC rescued the abrogated ppERK level in K-RasG12V-expressing cells (Fig. 4E). Together, our data suggest that C_16_H_20_FeClNO disrupts K-Ras/PM binding, nanoclustering and signaling through an ROS-mediated mechanism.

**Figure 4.**
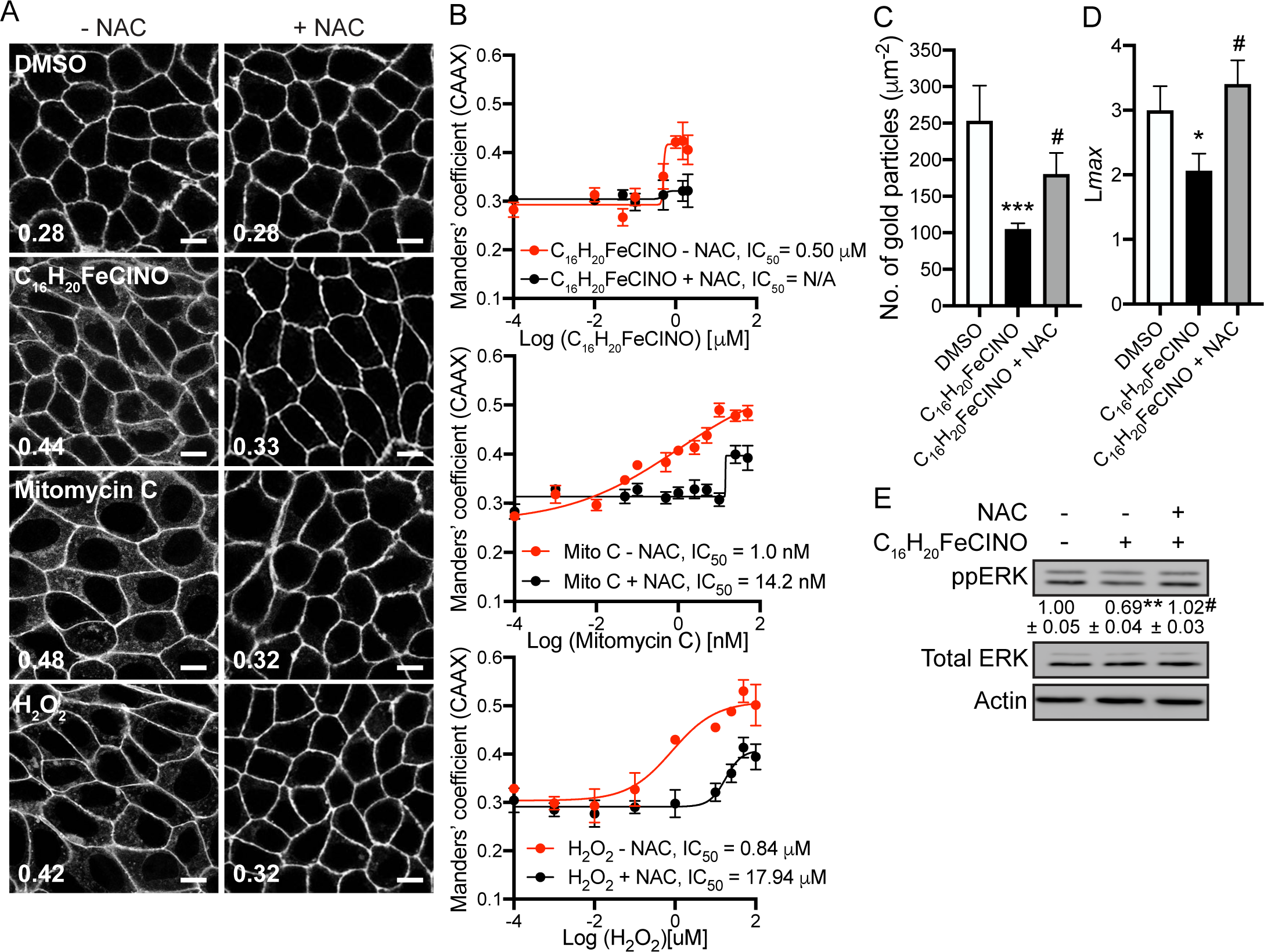
NAC supplementation returns K-Ras to the PM. (**A**) MDCK cells co-expressing GFP-K-RasG12V and mCherry-CAAX were treated with 2 μM C_16_H_20_FeClNO, 5 nM mitomycin C, 50 μM H_2_O_2_ in the presence or absence of 500 μM NAC for 48 h, and imaged by confocal microscopy. Representative images of GFP-RasG12V are shown. Inserted values represent an estimated mean fraction of mCherry-CAAX co-localized with GFP-RasG12V calculated by Manders’ coefficient from three independent experiments. Scale bar: 10 μm. (**B**) IC_50_s was estimated from the dose-response plots. (**C**) Basal PM sheets prepared from MDCK cells expressing GFP-K-RasG12V and treated with 2 μM C_16_H_20_FeClNO with or without 500 μM NAC for 48h were labeled with anti-GFP-conjugated gold and visualized by EM. Graphs show a mean number of gold particles ± S.E.M (n ≥ 15). (**D**) Spatial mapping of the same gold-labeled PM sheets was performed. The peak values, *L*_max_, of the respective weighted mean K-function *L(r) - r* curves are shown as bar graphs (*n* ≥ 15). (**E**) MDCK cells expressing GFP-K-RasG12V were treated with 1 μM C_16_H_20_FeClNO with or without 500 μM NAC for 48h. Cell lysates were blotted for phosphorylated ERK. Values indicate the mean ppERK ± S.E.M. from three independent experiments. Representative blots are shown. Total ERK and actin blots are shown as loading controls. Significant differences between control (DMSO-treated) and C_16_H_20_FeClNO-treated cells were assessed by one-way ANOVA tests for (**C**) and (**E**), and bootstrap tests for (**D**) (* p<0.05, ** p<0.01, *** p<0.001, # - not significant).

### ROS-induced K-Ras/PM dissociation is independent of PM PtdSer and K-Ras Ser181 phosphorylation

PtdSer is a phospholipid enriched in the inner leaflet of the PM and plays a critical role in the stable K-Ras/PM binding. K-Ras stably binds to the PM via the C-terminal farnesyl moiety in conjugation with the electrostatic interaction between the polybasic domain and the anionic head group of PtdSer (Cho *et al*., 2012b; Yeung *et al*., 2008; Zhou *et al*., 2017). Depletion of PM PtdSer content dissociates K-Ras from the PM and abrogates K-Ras signal output (Cho *et al*., 2012b; Cho *et al*., 2016b; Kattan *et al*., 2019; Kattan *et al*., 2021; Tan *et al*, 2018; van der Hoeven *et al*., 2018). Another mechanism regulating the stable K-Ras/PM binding is K-Ras phosphorylation at Ser181 by protein kinase C or G. Stimulated protein kinase C and G phosphorylate K-Ras Ser181 residue, located adjacent to the polybasic domain, which perturbs the electrostatic interaction of K-Ras and the negatively charged cellular membranes, resulting in K-Ras/PM dissociation (Bivona *et al*., 2006; Cho *et al*., 2016a; Kovar *et al*., 2020). To examine if C_16_H_20_FeClNO-induced K-Ras/PM dissociation is via these mechanisms, we studied cellular distribution of GFP-LactC2 (the C2 domain of lactadherin), a well-studied PtdSer probe (Yeung *et al*., 2008) and -K-RasG12V S181A, where Ser181 is substituted to Ala to prevent it from being phosphorylated. Our confocal microscopy shows that C_16_H_20_FeClNO treatment had minimal effects on the PM localization of LactC2 while K-RasG12V S181A redistributed from the PM (Fig. 5A), suggesting that C_16_H_20_FeClNO-induced K-Ras/PM dissociation is independent of PM PtdSer abundance and of K-Ras Ser181 phosphorylation.

**Figure 5.**
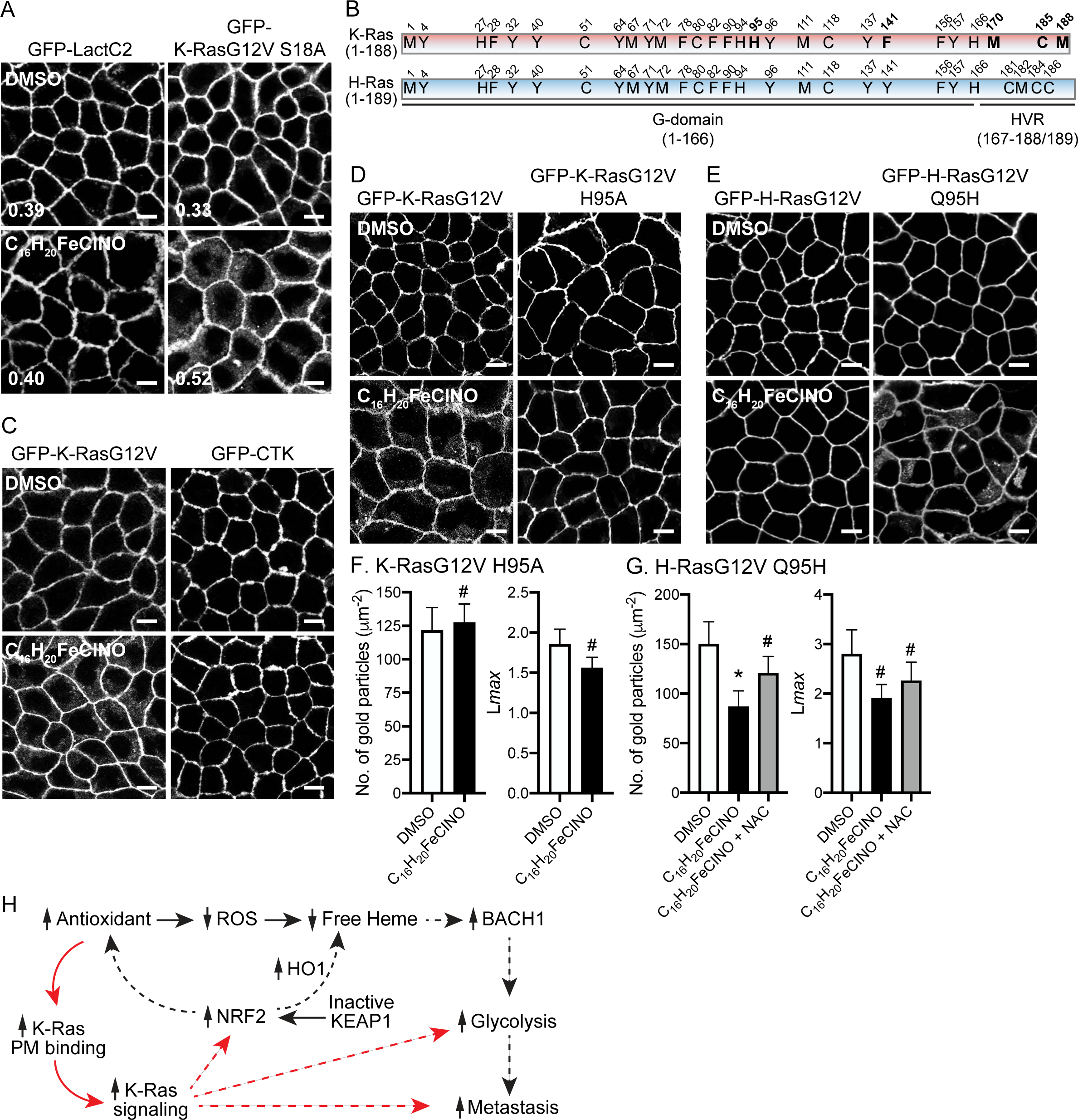
K-Ras His95 residue is involved in ROS-induced K-Ras PM dissociation. (**A**) MDCK cells co-expressing mCherry-CAAX and GFP-LactC2 or -K-RasG12V S181A were treated with 2 μM C_16_H_20_FeClNO for 48 h, and imaged by confocal microscopy. Inserted values represent an estimated mean fraction of mCherry-CAAX co-localized with GFP-tagged proteins calculated by Manders’ coefficient from three independent experiments. Scale bar: 10 μm. Representative images are shown. (**B**) A schematic diagram of K- and H-Ras amino acid residues that can be oxidatively modified. Amino acids unique to K-Ras are in bold. HVR – hypervariable region. MDCK cells stably expressing GFP-CTK (**C**), GFP-K-RasG12V or -K-RasG12V H95A (**D**), and GFP-H-RasG12V or -H-RasG12V Q95H (**E**) were treated with 2 μM C_16_H_20_FeClNO for 48 h, and imaged by confocal microscopy. Scale bar: 10 μm. Representative images from three independent experiments are shown. (**Left panels in F and G**) Basal PM sheets prepared from MDCK cells expressing GFP-K-RasG12V, -K-RasG12V H95A, -H-RasG12V, or -H-RasG12V Q95H and treated with 2 μM C_16_H_20_FeClNO with or without 500 μM NAC for 48h were labeled with anti-GFP-conjugated gold and visualized by EM. The graphs show a mean number of gold particles ± S.E.M (n ≥ 20). (**Right panels in F and G**) Spatial mapping of the same gold-labeled PM sheets was performed. The peak values, *L*_max_, of the respective weighted mean K-function *L(r) - r* curves are shown as bar graphs (*n* ≥ 20). Significant differences between control (DMSO-treated) and C_16_H_20_FeClNO-treated cells were assessed by Student’s tests for no. of gold particles and bootstrap tests for L*max*. (* p<0.05, # - not significant). **(H)** A model for the role of K-Ras in the antioxidant/BACH1/glycolysis-induced metastasis of K-Ras-driven NSCLC. Elevation of cellular antioxidant by antioxidant supplement or activating NRF2 blocks the oxidative modification of K-Ras His95 residue, enhancing K-Ras/PM binding and thereby K-Ras signaling. This in turn, stimulates Ras downstream effectors regulating glycolysis and metastasis, providing a positive feedback loop for the antioxidant/BACH1/glycolysis-induced metastasis of K-Ras-driven NSCLC. Solid and dotted lines indicate direct and indirect mechanisms, respectively. PM – plasma membrane, NFR2 - nuclear factor erythroid-derived 2-like 2, BACH1 - BTB and CNC homology 1, KEAP1 - Kelch-like ECH-associated protein 1, HO1 – heme oxidase-1.

### K-Ras His95 is critical for C_16_H_20_FeClNO-induced K-Ras/PM dissociation

While H-Ras is oxidatively modified perturbing its activity, little is known about K-Ras oxidative modifications (Burgoyne *et al*, 2012; Heo & Campbell, 2005; Heo *et al*, 2005; Messina *et al*, 2019). Since our data demonstrate that cellular ROS elevation disrupts the PM binding of K-Ras, but not H-Ras, which is reversed by NAC supplementation, we hypothesized that ROS oxidatively modifies K-Ras, blocking K-Ras/PM binding and its signaling. There are five amino acid residues that can be oxidatively modified and are unique to K-Ras (Fig. 5B). To identify the exact amino acid residue regulated by cellular ROS, we first generated MDCK cells stably expressing GFP-tagged <u>C</u>-terminal hypervariable region of <u>K</u>-Ras (GFP-CTK), sufficient for K-Ras transport to and stable interaction with the PM (Apolloni *et al*., 2000), and imaged its cellular localization after C_16_H_20_FeClNO treatment. We found that the compound had no effect on the PM localization of GFP-CTK (Fig. 5C), suggesting that the target residue must be located within the G-domain. In the G-domain, K-Ras contains His at the 95^th^ position that can be oxidatively modified (this position is Gln in H-Ras, which does not get oxidatively modified (Fig. 5B)). His95 is solvent exposed in the crystal structure of K-Ras (Fig. EV6), and previously reported molecular dynamics simulations suggest His95 would still be accessible to solvent, and hence to ROS when K-Ras is bound to a model membrane (Prakash *et al*, 2019; Prakash *et al*, 2016). To examine if His95 is involved in the ROS-induced K-Ras/PM dissociation, we substituted the His with Ala, which does not undergo oxidative modification. MDCK cells expressing GFP-K-RasG12V H95A were treated with C_16_H_20_FeClNO, and K-Ras cellular localization was studied by confocal and electron microscopy. C_16_H_20_FeClNO treatment had no effect on the PM binding and nanoclustering of K-RasG12V H95A (Figs. 5D and F). We further substituted the Gln95 of H-Ras to His and studied its cellular localization after C_16_H_20_FeClNO treatment. While Gln95His mutation did not alter the PM binding and nanoclustering of H-RasG12V under basal conditions (Fig. EV7), C_16_H_20_FeClNO treatment abrogated the PM binding and nanoclustering of H-Ras G12V Q95H, and this effect was reversed by NAC supplementation (Figs. 5E and G). Together, our data suggest that cellular ROS regulates the PM binding and nanoclustering of K-Ras by oxidative modification at His95 residue.

In this study, we have demonstrated that C_16_H_20_FeClNO translocates PM K-Ras to the cytosol and endomembranes by elevating cellular ROS, and it blocks K-Ras signaling and the growth of K-Ras-dependent PDAC and NSCLC cells. These effects were reversed by antioxidant supplementation. While previous studies have reported the roles of K-Ras signaling on regulating redox balance, our study reveals direct effects of the redox balance on K-Ras/PM binding and its signal output, thus providing a mechanistic link between K-Ras and recent reports on the antioxidant/BACH1/glycolysis-promoted metastasis of K-Ras-driven NSCLC (Lignitto *et al*., 2019; Wiel *et al*., 2019). Under oxidative stress, the oxidation of heme-containing proteins releases free heme, which stimulates degradation of BACH1 (BTB and CNC homology 1), a pro-metastatic transcriptional factor (Lignitto *et al*., 2019; Zenke-Kawasaki *et al*, 2007). Prolonged supplementation of antioxidants reduces cellular ROS levels, which lowers cellular free heme, stabilizing BACH1 protein. The elevated BACH1 further promotes glycolysis by upregulating transcription of *GAPDH* and *hexokinase-2*, promoting metastasis in K-Ras-driven NSCLC (Wiel *et al*., 2019) (Fig. 5H). Consistently, promoting the endogenous antioxidant program either by activating NRF2 (nuclear factor erythroid-derived 2-like 2), the master transcriptional regulator of antioxidant responses, or blocking its negative regulator, KEAP1 (Kelch-like ECH-associated protein 1), upregulates heme oxygenease-1, which degrade free heme, resulting in the BACH1-dependent metastasis of K-Ras-driven NSCLC (Li & Stocker, 2009; Wiel *et al*., 2019). These studies have thus demonstrated the role of antioxidants in the BACH1/glycolysis-mediated metastasis of K-Ras-driven NSCLC, and our study adds the pivotal role of oncogenic K-Ras in this signal axis. We propose that in K-Ras-driven NSCLC, elevation of cellular antioxidants prevents the oxidative modification of the His95 residue of oncogenic mutant K-Ras, enhancing K-Ras/PM binding, and thereby K-Ras signal output. This elevated K-Ras signaling further promotes NRF2 expression, stimulating the NRF2-mediated antioxidant program (DeNicola *et al*, 2011; Singh *et al*, 2008; Yang *et al*, 2021). This in turn, provides a positive-feedback loop for the antioxidant/BACH1/glycolysis-induced metastatic signaling. The enhanced oncogenic K-Ras signaling also promotes glycolysis and metastasis pathways (Cox *et al*., 2014; Gorfe & Cho, 2019; Ying *et al*, 2012), further promoting the antioxidant/BACH1/glycolysis-induced NSCLC metastasis (Fig. 5H).

We have identified K-Ras His95 as a novel amino acid residue for regulating K-Ras/PM binding via oxidative modification. Although the exact mechanisms on how oxidative modification of K-Ras His95 perturbs the PM binding needs to be further elucidated, the observations that ROS elevation does not affect the PM binding of K-Ras4A, which also has His95, and that Gln95His substitution dissociates H-Ras from the PM (Figs. 2 and 5) provide a hint into the possible involvement of the PDE6δ-Arl2/3 transport machinery. When K-Ras and depalmitoylated H- and N-Ras are dissociated from the PM, PDE6δ continuously sequesters them from endomembranes via interacting with their C-terminal farnesyl moiety. The release factors, Arl2 and 3 then bind the Ras/PDE6δ complex and release Ras to perinuclear membranes for returning to the PM (Chandra *et al*., 2012; Ismail *et al*, 2011; Schmick *et al*., 2014). While PDE6δ does not bind depalmitoylated K-Ras4A likely due to steric hindrance caused by two Lys residues immediately prior to the C-terminal farnesylated Cys of K-Ras4A, PDE6δ directly interacts with five amino acids (residues 180-184) immediately prior to the farnesylated Cys of K-Ras for the stable binding (Dharmaiah *et al*, 2016). Thus, it is plausible that oxidatively modified His95 of K-Ras may perturb Arl2/3 interaction with the K-Ras/PDE6δ complex, resulting in K-Ras distribution from the PM to endomembranes. In conclusion, our study demonstrate that redox system directly regulates K-Ras/PM binding and its signaling via oxidative modification at His95, and proposes a role of oncogenic mutant K-Ras in the antioxidant/BACH1/glycolysis-induced metastasis signaling pathway in K-Ras-driven NSCLC.

## Methods and Materials

### Synthesis of 1-ferrocenyl-2-[(dimethylamino)methyl]-2-propen-1-one

(Compound **1**). All synthetic manipulations were done under a nitrogen atmosphere unless otherwise noted. All glassware were oven dried at 110 °C for 12 hours before use. Acetylferrocene was prepared according to the literature procedure (Donahue & Donahue, 2013; Graham *et al*, 1957). Bis(dimethylamino)methane, phosphoric acid, acetic acid, and hydrochloric acid (1M) in ether were purchased from commercial sources and used as received. Solvents were dried with a solvent purification system from Inert Pure Company (THF, CH_2_Cl_2_, Et_2_O, and toluene) and degassed using three freeze-pump-thaw cycles before use. All solvents were stored over 4 Å molecular sieves in a nitrogen-filled glove box. CDCl_3_ (99.9%) and D_2_O (99.9%) were purchased from Acros Laboratories and dried over 4 Å molecular sieves before use. ^1^H and ^13^C NMR spectra were recorded on a Bruker 300 MHz NMR spectrometer. Spectra were referenced to the residual solvent as an internal standard for ^1^H NMR: CDCl_3_, 7.26 ppm, D_2_O 4.79 ppm; for ^13^C NMR: CDCl_3_, 77.16. Coupling constants (J) are expressed in hertz (Hz). Elemental analyses were performed by Midwest Microlab, LLC, in Indianapolis, IN.

Acetylferrocene (1.39 g, 6.1 mmol) was added to a premixed solution of bis(dimethylamino)methane (1.47 mL, 1.77 mmol), phosphoric acid (0.65 mL, 9.5 mmol) and 14 mL of acetic acid in a two-neck round bottom flask equipped with a stir bar and condenser. Under a nitrogen atmosphere, the mixture was refluxed at 110 °C and stirred for 5 hours. The mixture was allowed to cool to room temperature, diluted with 14 mL of water, and extracted two times with 50 mL of diethyl ether. The aqueous solution was then cooled using an ice-water mixture and made alkaline by adding solid NaOH pellets (6 g) and extracted three times with 50 mL of diethyl ether. Finally, the organic solution was washed with water, dried in MgSO_4,_ and concentrated under reduced pressure. The crude residue was passed through short silica gel column chromatography using (Hexane:DCM:Et_3_N, 80:10:10) and yielded 1-ferrocenyl-2-[(dimethylamino)methyl]-2-propen-1-one. 1.29 g, 71% yield. ^1^H NMR (CDCl_3_, 300 MHz): d 5.86 (s, 1H), 5.69 (s, 1H), 4.84 (s, 2H), 4.51 (s, 2H), 4.24 (s, 5H), 3.25 (s, 2H), 2.28 (s, 6H). ^13^C NMR (CDCl_3_, 75 MHz): d 200.1, 146.9, 121.7, 77.9, 72.2, 70.9, 70.1, 61.6, 45.5.

### Synthesis of 1-ferrocenyl-2-[(dimethylamino)methyl]-2-propen-1-one hydrochloride

(Compound **2**). Compound **1** (1.19 g, 4.0 mmol) and dichloromethane (20 mL) was charged into a 40 mL glass vial, and the reaction mixture was cooled using an ice bath. Hydrochloric acid (1M solution in diethyl ether, 4 mL, 4.0 mmol) was added dropwise. Reaction contents were stirred for 1 hour, and the solvent was removed under vacuum; the residue was washed with diethyl ether (2 x 10 mL) and dried under vacuum, yielding compound **2**, 1.27 g, 95% yield. ^1^H NMR (D_2_O, 300 MHz): d 6.73 (s, 1H), 6.37 (s, 1H), 4.93-4.84 (m, 4 H), 4.31 (s, 5 H), 3.98 (s, 2 H), 2.87 (s, 2H). ^13^C NMR (D_2_O, 75 MHz): d 200.3, 136.3, 134.98, 75.6, 74.63, 71.52, 70.85, 70.57, 59.47, 42.64. HRMS (ESI) for [C_16_H_20_FeNO]^+^[M]^+^ Calcd. 298.0889 Found. 298.0884. Anal. Calcd. For: C_16_H_20_ClFeNO: C, 57.60, H, 6.04, N, 4.20; Found: C, 56.87, H, 5.89, N, 4.21.

### Plasmids and Reagents

The following antibodies to measure Ras signaling were purchased from Cell Signaling Technology (Danvers, MA): pAkt ((Ser473) (D9E) XP; Cat #4060L), total Akt ((pan) (40D4); Cat # 2920S), ppErk (p44/42 MAPK (Erk1/2) (Thr202/Tyr204) (D13.14.4E)) XP; Cat #4370L), total Erk (p44/42 MAPK (Erk1/2) (L34F12). The following antibodies used to measure housekeeping genes were purchased from Proteintech (Rosemont, IL): β-actin (Cat # 66009-1-Ig) and GFP-tag (Cat # 60002-1-Ig). Forward and reverse primers for GFP-K-RasG12V H95A, GFP-K-RasG12V H95C and GFP-H-RasG12V Q95H mutants were designed and ordered from Agilent QuikChange Primer Design (Agilent; Santa Clara, CA). cDNA of H-RasG12V Q95H, K-RasG12V H95A and H95C were cloned into a pBLST vector. Correct mutation and sequence alignment was ensured by Sanger Sequencing by Genewiz (South Plainfield, NJ) and Basic Local Alignment Search Tool (BLAST; NIH National Library of Medicine; Bethesda, MD). The RFP-tagged organelle markers were purchased from Invitrogen (Carlsbad, CA): CellLight Golgi-RFP BacMam 2.0 (Cat# C10593), CellLight Lysosomes-RFP BacMam 2.0 (Cat # C10597), CellLight Mitochondria-RFP BacMam 2.0 (Cat # C10601), CellLight Late Endosome-RFP BacMam 2.0 (Cat #C10589), CellLight Early Endosome-RFP BacMam 2.0 (Cat #C10587), and ER-Tracker Red ((BODIPY TR Glibenclamide) (Cat #E34250)). Mitomycin C was purchased from Alfa Aesar (Cat # J63193). Carmustine was purchased from MedChemExpres (Cat # HY-13585/CS-2935). N-acetylcysteine was purchased from Sigma-Aldrich (Cat # A7250-10G). Hydrogen peroxide (H_2_O_2_) was purchased from Sigma Aldrich (Cat #H1009).

### Cell Culture

Madin-Darby Kidney Cells (MDCK) were maintained in Dulbecco’s modified eagle medium (DMEM; Gibco; Cat #10569-010). Human pancreatic ductal adenocarcinoma cells (PDACs): BxPC3, Panc 10.05 and AsPC-1 were maintained in RPMI-1640 (ATCC; 30-2001), MiaPaCa2 and PANC-1 were maintained in DMEM, HPAC were maintained in DMEM-F12 (Gibco; Cat # 11320033) and HPAF-II were maintained in EMEM (ATTC; 30-2003). Human non-small cell lung cancer cell (NSCLCs): H522, H1975, H1299, A549, H23, H441, H1703 and H358 were maintained in RPMI-1640. All cancer cell lines were maintained in media supplemented with 10% Fetal Bovine Serum (Gibco; Cat #16000-069) and 2mM L-glutamine (GenDEPOT; Cat # CA009-010). Cells were test for mycoplasma (MycoAlert PLUS Mycoplasma Detection Kit; Cat# LT07-710). All cell lines were maintained in an incubator at 37°C at 5% CO2.

### Proliferation Assay

Pancreatic ductal adenocarcinoma cells were seeded at 3x10^5^ and non-small cell lung cancer cells were seeded at 1x10^5^-2x10^5^ on a 96-well plate. 24 hours later, cells were incubated with DMSO (control) or increasing concentrations of C_16_H_20_FeClNO in 100 μL complete growth media for 72 hours, changing the media every 24 hours. Cell proliferation was quantified using CyQuant NF Cell Proliferation Assay Kit (Molecule Probes; Cat # 35006) according to the manufacturer’s protocol. Plates were read using BioTek Synergy H1 microplate reader at excitation/emission of 480/530 nm.

### Confocal Microscopy

MDCK cells were seeded on coverslips at 2.50x10^5^ on a 12-well plate. Cells were treated with increasing concentrations of C_16_H_20_FeClNO, mitomycin C, carmustine or H_2_O_2_ for 48 hours. Cells were washed twice with ice-cold 1x phosphate-buffered saline (PBS) and then fixed with 4% paraformaldehyde (Electron Microscopy Services, Cat #15710)) followed by 50 mM NH_4_Cl. Slides were imaged using the Olympus FV1000 confocal microscope using a 60x objective. Images were quantified for co-localization using Manders’ coefficient on ImageJ software (version 1.52a).

### Electron microscopy (EM) and spatial mapping

Plasma membrane sheets were prepared and fixed as previously described (Garrido *et al*., 2020; Hancock & Prior, 2005; Prior *et al*., 2003). For univariate analysis, plasma membrane sheets were labeled with anti-green fluorescent protein (anti-GFP) antibody conjugated to 4.5-nm gold particles. Digital images of the immunogold-labeled plasma membrane sheets were taken in a transmission electron microscope. Intact 1 μm^2^ areas of the plasma membrane sheet were identified using ImageJ software, and the (*x*, *y*) coordinates of the gold particles were determined (Hancock & Prior, 2005; Prior *et al*., 2003). K-functions (Ripley, 1977) were calculated and standardized on the 99% confidence interval (CI) for univariate functions (Diggle, 2000; Hancock & Prior, 2005; Prior *et al*., 2003). In the case of univariate functions, a value of *L*(*r*) - *r* greater than the CI indicates significant clustering, and the maximum value of the function (*L*_max_) estimates the extent of clustering. Differences between replicated point patterns were analyzed by constructing bootstrap tests as described previously (Diggle, 2000; Plowman *et al*, 2005), and the statistical significance against the results obtained with 1,000 bootstrap samples was evaluated.

### Subcellular Fractionation Assay

MDCK cells were seeded at 17x10^5^ onto 10cm dishes. Then, cells were treated with increasing concentrations of C_16_H_20_FeClNO, mitomycin C, carmustine or H_2_O_2_ for 48 hours. Dishes were washed twice with ice-cold 1x PBS. Lysates were harvested in buffer A containing 10mM Tris pH 7.5, 75mM NaCl, 25mM NaF, 5mM MgCl_2_, 1mM EGTA, 1M DTT and 100 μM NaVO_4_. Lysates were spun at 100,000xg for 1 hour at 4°C using Sorvall Discovery MX120SE Ultracentrifuge (Thermo Scientific; Waltham; MA) to isolate the cytosolic fraction. Pellets were resuspended in Buffer A and sonicated 10x at 4°C to isolate the membrane-bound fraction. Protein concentrations were determined using the BCA protein assay (Thermo Fisher Scientific; Reagent A Cat #23228; Reagent B Cat # 1859078). SDS-page and immunoblotting were performed using 20-25 μg of lysate from each sample group. Blots were detected using chemiluminescence on the Amersham Imager 600 (GE Healthcare Life Sciences; Marlborough, MA). Blots were quantified using ImageJ software.

### Western Blotting

MDCK cells were seeded at 3x10^5^ on a 6-well plate and treated with increasing concentrations of C_16_H_20_FeClNO for 48 hours. Then, cells were washed twice with ice-cold 1x PBS. Cells were harvested in a lysis buffer consisting of 50mM Tris pH 7.5, 25 mM NaF, 5 mM MgCl_2_, 75 mM NaCl, 5 mM EGTA, 1 mM DTT, 100 μM NaVO_4_ and 1% NP-40.

Protein concentrations were determined using the BCA protein assay. SDS-page and immunoblotting were performed using 20-25 μg of lysate from each sample group. Blots were detected using chemiluminescence on the Amersham Imager 600. Blots were quantified using ImageJ software.

### Quantification of ROS levels by H_2_DCFDA

Wild-type MDCK cells were seeded at 2 x 10^4^ on a clear-bottomed 96-well plate overnight. Then, cells were co-treated with 25 μM of DCFDA dye (H_2_DCFDA; Thermo Fisher Scientific; Cat # D399) and either H_2_O_2_, C_16_H_20_FeClNO only or C_16_H_20_FeClNO with NAC for 48 hours. After 48 hours, fluorescence was read using the BioTek Synergy H1 microplate reader with emission/excitation at 495/527nm.

### Statistics

GraphPad prism software (v.8.4.0) was used for one-way ANOVA and unpaired *t*-test.

## Acknowledgements

This work was supported by NIH [R01 GM144836] to A.A.G, Wright State Seed Grant Program to K.-J.C., and NIH [R15 CA232765], The American Chemical Society Petroleum Research Fund [PRF-59893-UR7] and Wright State University CoSM fund to K.A.

## Author Contributions

K.M.R. and K.-J.C. designed and performed the experiments, analyzed the data, and wrote the manuscript text. J.S. and D.H. prepared the ferrocene compound and S.A. provided Fig. 1A and EV1. A.A.G. provided K-Ras crystal structure in Fig. EV6 and participated in data discussion. All authors reviewed the manuscript.

---

**Figure EV1.**
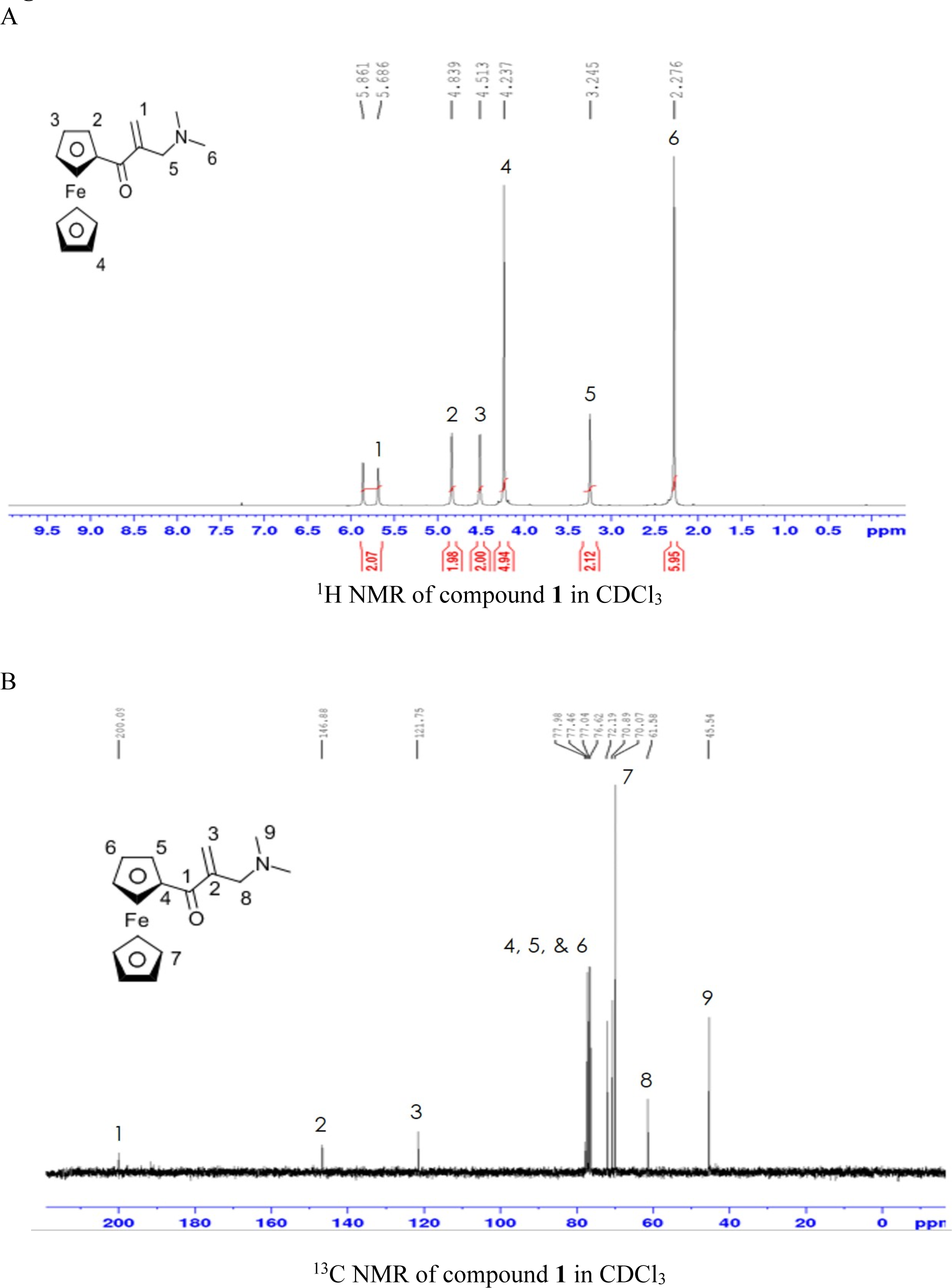
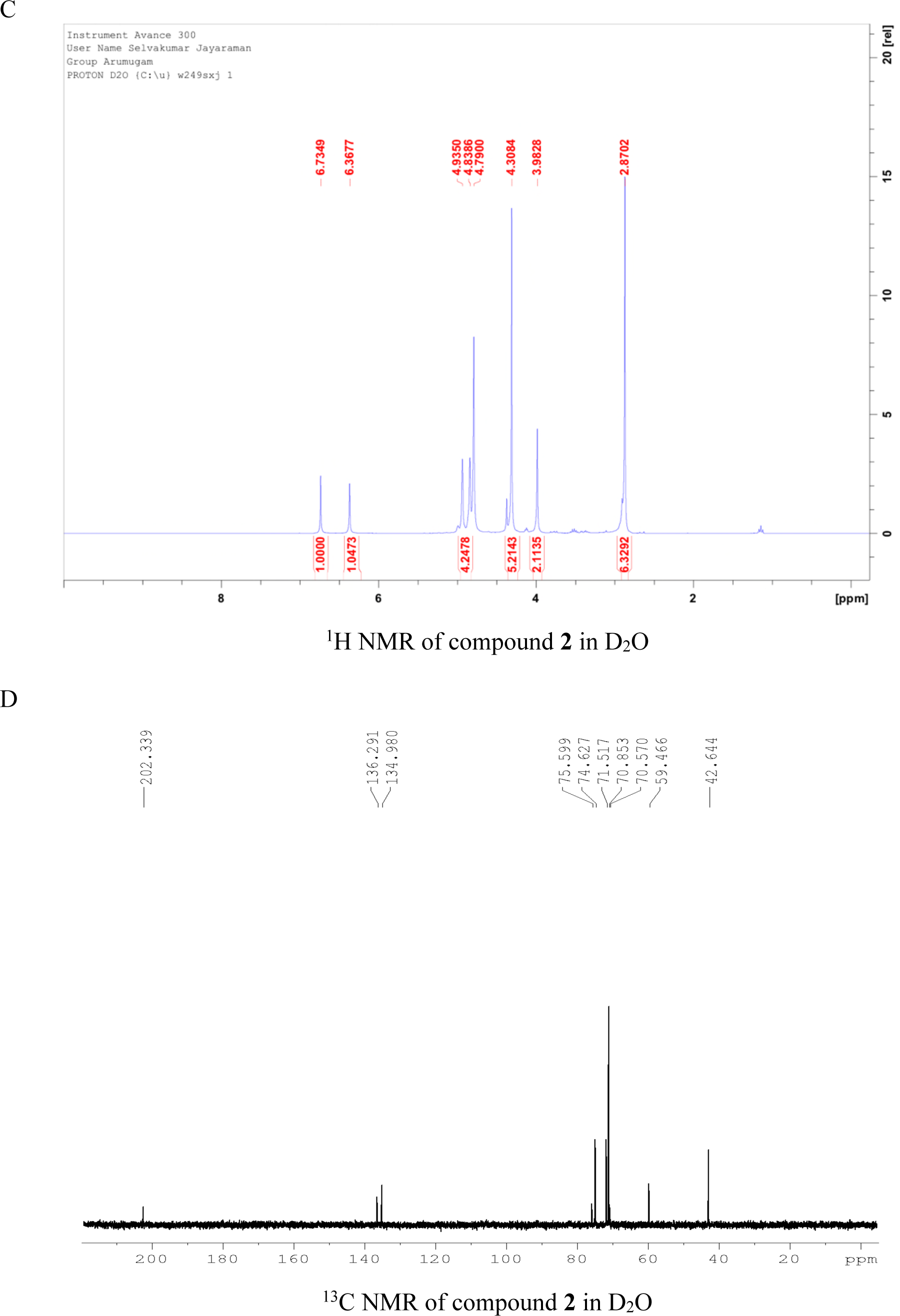

**Figure EV2.**
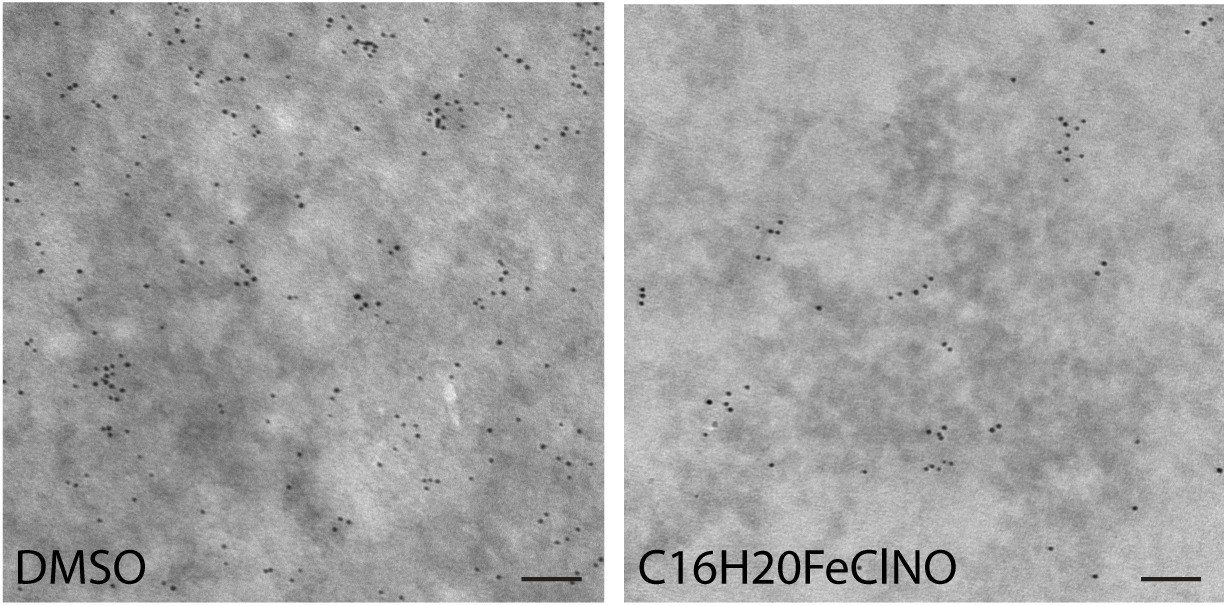
Basal PM sheets prepared from MDCK cells expressing GFP-K-RasG12V and treated with DMSO or 2 μM C_16_H_20_FeClNO for 48h were labeled with anti-GFP-conjugated gold and visualized by a transmission EM. Representative images are shown from n ≥ 20. Each dot represents GFP-K-Ras localized to the plasma membrane. Scale bar: 100 nm.

**Figure EV3.**
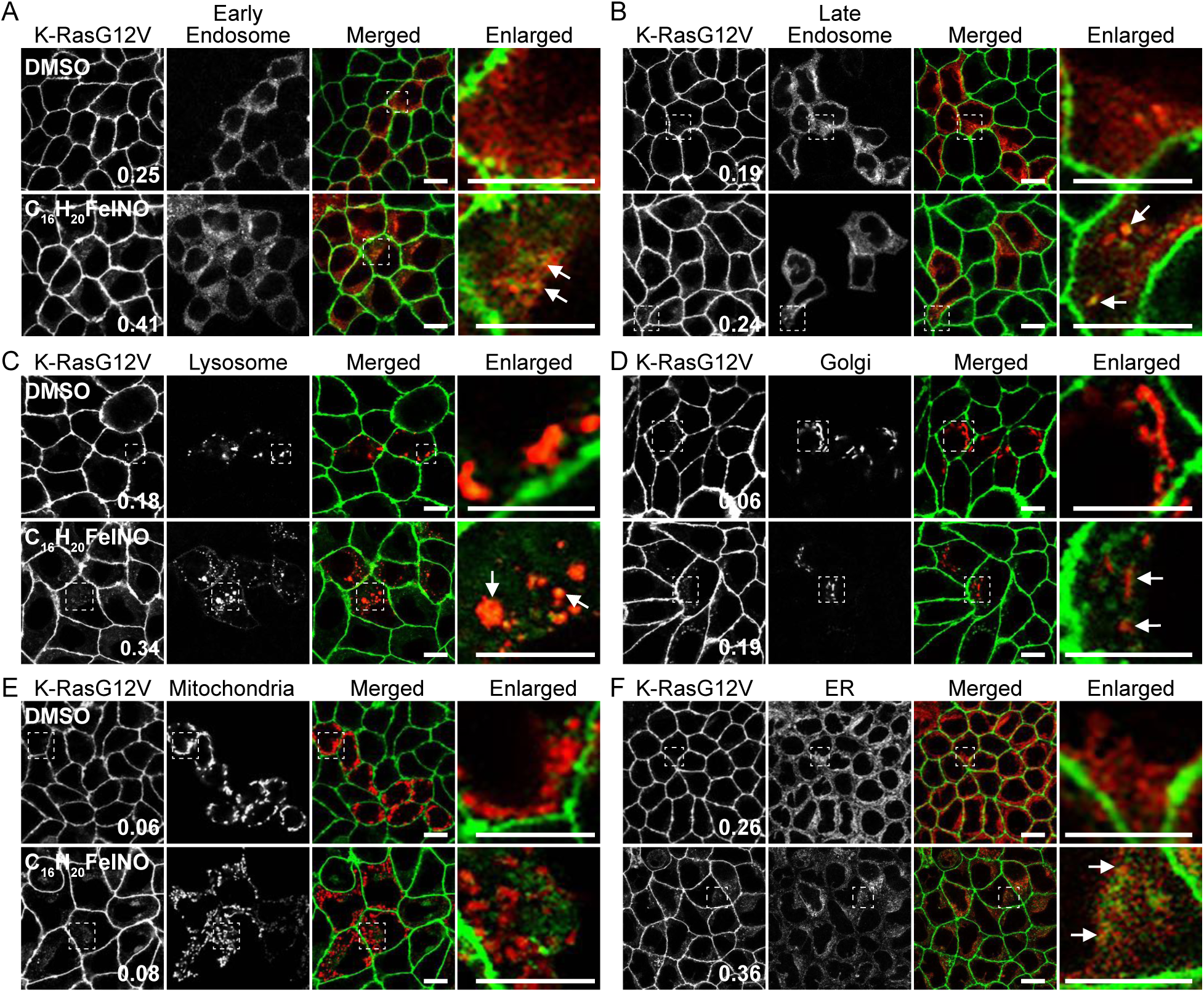
MDCK cells stably expressing GFP-K-RasG12V were incubated with 2 μM C_16_H_20_FeClNO for 48 in the presence of modified baculovirus encoding RFP-tagged organelle markers (**A – E**), or 2 μM ER-tracker for the final 1 h of incubation in C H FeClNO (**F**). Cells were fixed with 4% PFA and imaged in a confocal microscope. Selected regions indicated by the white squares are shown at a higher magnification. K-RasG12V that is co-localized with the markers is indicated by arrowheads. Scale bar: 10 μm. Values indicate K-RasG12V co-localized with the markers (indicated by arrowheads) calculated by Manders’ coefficients from n ≥ 15 images.

**Figure EV4.**
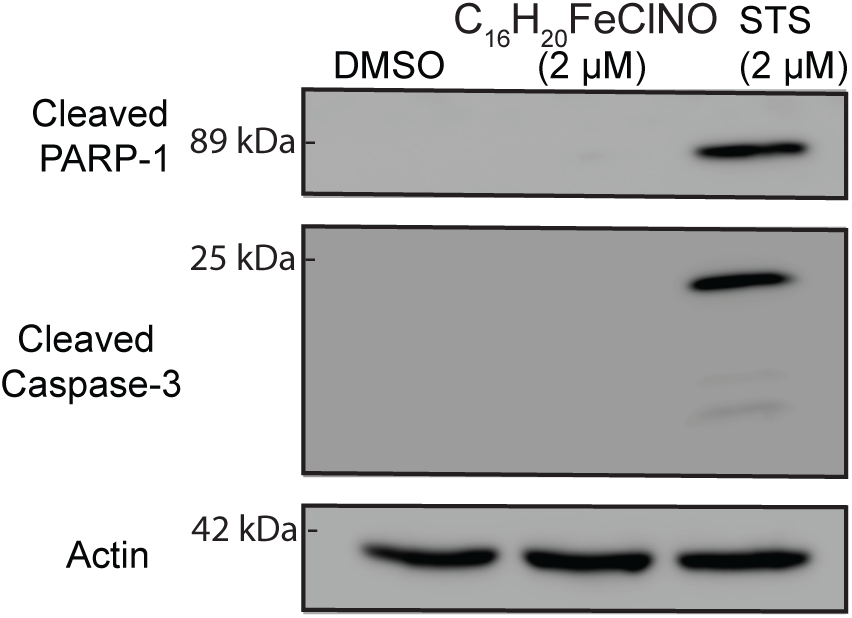
MDCK cells expressing GFP-K-RasG12V were treated with DMSO, 2 μM C_16_H_20_FeClNO for 48h or 2 μM staurosporine (STS) for 6h. Cell lysates were blotted for cleaved PARP-1 and cleaved caspase-3. Representative blots are shown from three independent experiments. An actin blot is shown as a loading control.

**Figure EV5.**
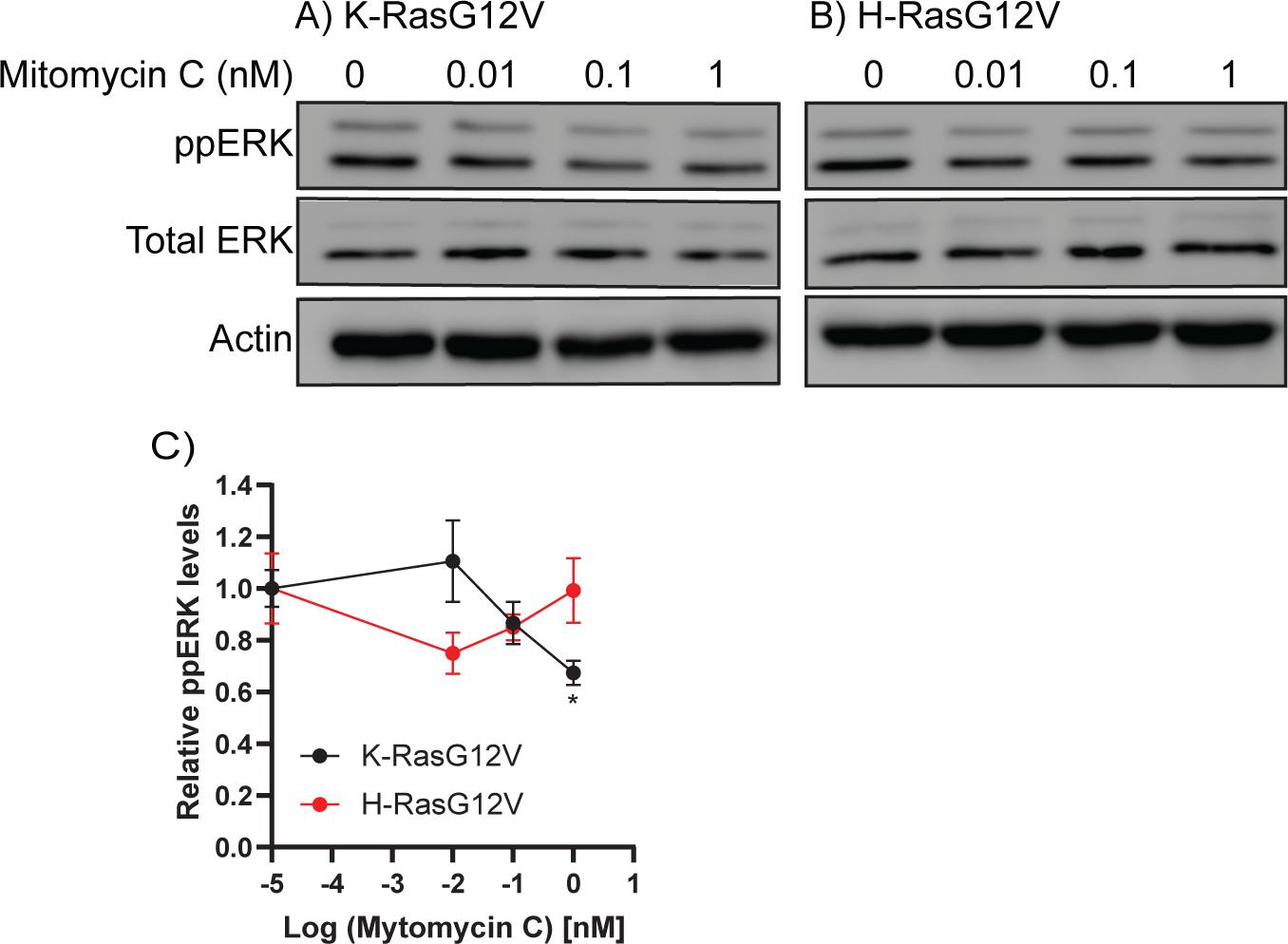

**Figure EV6.**
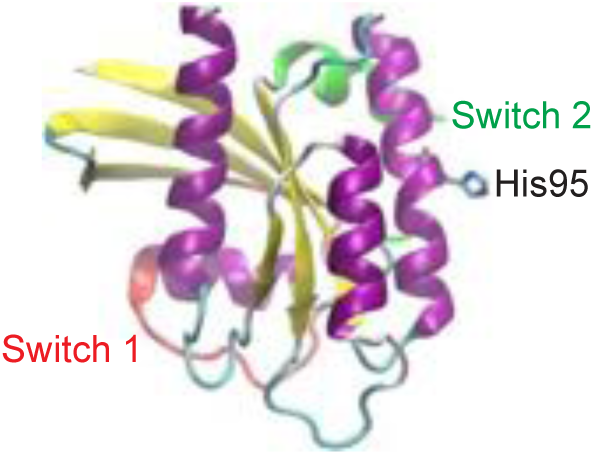
The K-Ras catalytic domain structure shows that His95 side chain is solvent exposed and thus available for oxidative modification.

**Figure EV7.**
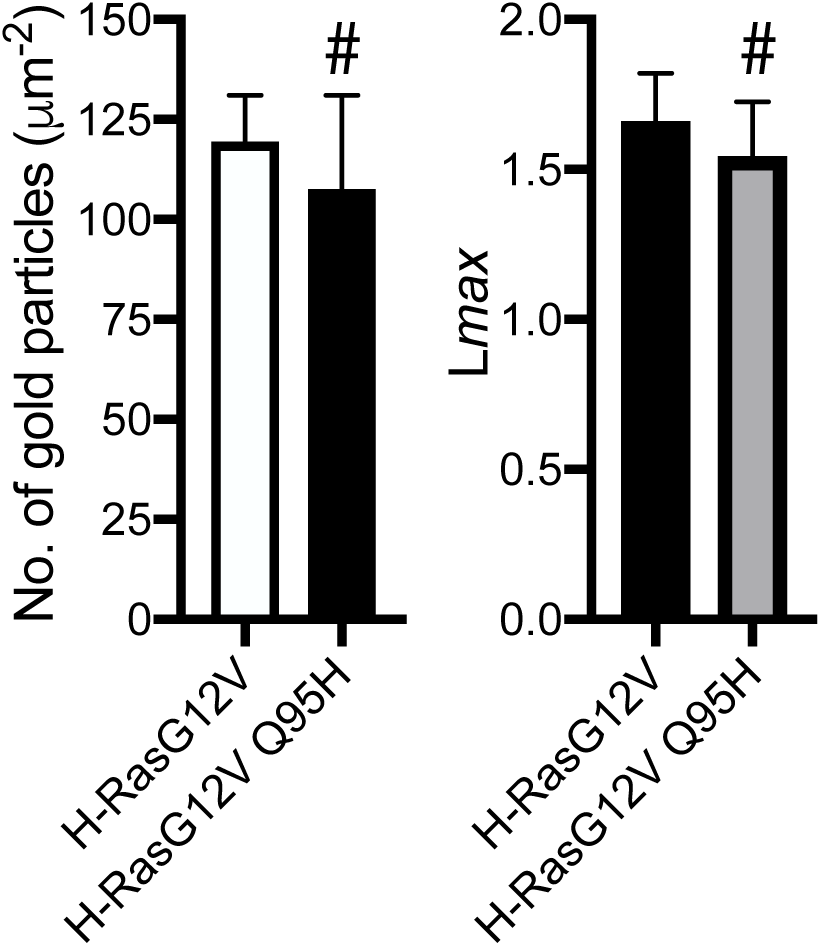
(**Left panel**) Basal PM sheets prepared from MDCK cells expressing GFP-H-RasG12V or -H-RasG12V Q95H were labeled with anti-GFP-conjugated gold and visualized by EM. The graph shows a mean number of gold particles ± S.E.M (n ≥ 20). (**Right panel**) Spatial mapping of the same gold-labeled PM sheets was performed. The peak values, *L*_max_, of the respective weighted mean K-function *L(r) - r* curves are shown as bar graphs (*n* ≥ 20). Significant differences between H-RasG12V and H-RasG12V Q95H were assessed by Student’s *t*-tests for (left panel) and bootstrap tests for (right panel) (# - not significant).

